# Novel plant-derived bioactive compounds shape the human gut microbiome *in vitro*

**DOI:** 10.1101/2024.05.07.592791

**Authors:** Karla E Flores Martinez, Clay S. Bloszies, Bethany M. Henrick, Steven A Frese

## Abstract

Mounting evidence supports the potential of dietary bioactive compounds to reduce chronic disease risk. Recently, the biological activity of N-trans-caffeoyltyramine (NCT) and N-trans-feruloyltyramine (NFT) has been hypothesized to drive regulation of gut permeability,but the impact of these components on the human gut microbiome composition has not been studied. The aim of this work is to determine whether purified NCT and NFT, or a hemp hull product containing NCT and NFT (Brightseed® Bio Gut Fiber), can impact the gut microbiome using an *in vitro* fermentation assay. To address this question, we treated three fecal inocula, representative of the human gut microbiome, with Bio Gut Fiber™ and NCT/NFT and evaluated their respective impact against starch and methylcellulose as controls. We found strong changes exerted by Bio Gut Fiber™ and NCT/NFT on the gut microbiome relative to starch and methylcellulose, with distinct responses across all microbial communities. Among communities treated with Bio Gut Fiber™, we saw increased community productivity and increased community diversity. Further, to determine whether changes found in gut microbiome profiles were dose-dependent, we tested different concentrations of NCT/NFT and found a dose-dependent impact on the resulting microbial compositions. Through this work, we provide novel insight into the potential of bioactive components to shape the gut microbiome, highlighting the potential for plant-derived bioactives to improve the human gut microbiome and host health.

## 1. Introduction

Over the past decade, understanding the considerable impact of diet on health and disease has increased interest in developing nutritional strategies to prevent a broad range of morbidities^1^. Various studies have evidenced plant-derived bioactive components as a potential approach for minimizing the risk of developing chronic diseases^2^. Bioactives are defined as “naturally occurring compounds with biological activity that have evidenced a beneficial effect on human health beyond conventional nutrition.”. Bioactive components are present in grains, vegetables, fruits, herbs, and other plant sources and include carotenoids, polyphenols, vitamins, bioactive peptides, and dietary fiber^2^.

A growing body of work has associated plant-derived bioactive compounds with modulation of the gut microbiome^2,3,4^. The gut microbiome is home to a diverse community of microbes that inhabits the gastrointestinal tract and the interaction of these microbes with dietary components can significantly affect host health^3^. In particular, dietary fiber has demonstrated prebiotic activity with significant effects on the growth and activity of beneficial gut microbiota^5^. Fibers such as galacto-oligosaccharides (GOS), fructo-oligosaccharides (FOS), and resistant starch (RS) reach the colon and are fermented by bacteria into short-chain fatty acids (SCFAs)^5^ that can strengthen gut barrier function and mitigate enteric inflammation^6,7^. Consequently, there is growing interest in understanding the interactions between complex dietary fiber and the gut microbiome, but most studies to date have focused on dietary fiber available through whole foods^8,9^ or structurally repetitive fibers such as GOS and FOS^10,11,12^.

Hemp (*Cannabis sativa L*.) is composed of 25-30% oil and 25-30% protein contained in the hemp seed, while the exterior layer, known as the hull, contains 30-40% fiber^8,13^. Despite the fiber fraction comprising one-third of the composition of hemp, most of the research has mainly focused on the oil and protein fractions of the hemp seed^8^. Aside from its macronutrient composition, hemp seed hull contains bioactive compounds such as *N*-trans caffeoyltyramine (NCT) and *N*-trans-feruloyltyramine (NFT), and these compounds are thought to play a protective role in gut barrier function and reduce inflammation^13^. However, there is little current knowledge as to whether or how NCT and NFT or hemp hull fiber can impact gut microbiome composition and function.

Here, we examined the impact of a novel hemp hull ingredient (Brightseed® Bio Gut Fiber) as well as NCT/NFT in an *in vitro* fermentation model using inocula representative of three deeply phenotyped human gut microbiomes. In addition to these hemp-derived ingredients, we compared these three communities on an insoluble fiber (methylcellulose) and starch as control substrates. Using 16S rRNA amplicon sequencing, we compared the resulting communities and assessed the impact of Bio Gut Fiber™, NCT/NFT, and methylcellulose on human gut microbiomes *in vitro*, in comparison to a starch control medium. We also assessed community productivity through growth kinetics and short-chain fatty acid (SCFA) quantification.

## 2. Methods

### 2.1 In vitro batch fermentation

Fecal samples were collected from healthy individuals under the supervision of the University of Nevada, Reno Institutional Review Board (Approval #1751022), from which multiple aliquots were collected and stored at - 80°C. Three standardized fecal inocula were prepared by pooling fecal samples (N = 6 individuals per inoculum) profiled by 16S rRNA amplicon sequencing and identified to belong to one of three distinct compositional groups^14^ to generate three distinct inocula: E1, E2, and E3. To prepare the inocula, frozen samples were diluted 1:10 (w/v) in ice-cold PBS (pH 7.0) containing 15% glycerol.

Each inoculum was added (1% v/v) to a 96-well deep well plate containing 1 mL of anaerobic cultivation medium adapted from Aranda-Díaz *et al*.^15^, supplemented with soluble starch (0.35% w/v) which is included in the adapted culture medium recipe as an added complex carbohydrate source, or, in place of the starch, methylcellulose (2% w/v), Bio Gut Fiber™ (BGF; 2% w/v), or NCT/NFT (2% w/v). Six independent replicates per enterotype and treatment were used. BGF was derived from hemp hulls (*Cannabis sativa L*.), and NCT/NFT were obtained from Brightseed, Inc. (South San Francisco, California, USA).

The anaerobic culture medium used included BHI medium supplemented with 0.3 g/L of L-cysteine HCl, 0.3 g/L of sodium thioglycolate, 1.5 mg/L of vitamin K1, and 0.3 mg/L of hemin, described here as mBHI. mBHI contains 0.2% glucose as an ingredient to support minimal levels of carbohydrate-dependent growth. After inoculation, 100 µL of each replicate was transferred to a 96-well untreated cell culture plate and sealed with a sterile, gas-permeable sealing film (Diversified Biotech, Dedham, MA, USA) and incubated at 37°C in a Cerillo Stratus Kinetic 96-well plate reader (Cerillo, Charlottesville, VA, USA) with constant shaking and measuring optical density at 600 nm (OD_600nm_) every 3 mins for 24 hours. The 96-well deep well plate was sealed with a sterile, breathable sealing film (Celltreat Scientific Products; Pepperell, MA, USA) and incubated at 37°C for 24 hours. Both the 96-well plate and the 96-well deep well plate were incubated anaerobically (90% N_2_, 5% CO_2_, 5% H_2_) at 37°C in an anaerobic chamber (Coy Laboratory Products, Grass Lake, MI, USA) with a digital oxygen sensor and palladium catalysts to passively remove any residual oxygen in the chamber. After 24 hours, the 96-well deep well plate was removed and centrifuged at 4°C for 20 minutes (4,000 RPM). From each well, 500 µL of cell-free supernatant was collected, transferred into a new 96-well deep well plate, and stored at -80°C. The pelleted cells were resuspended in DNA/RNA Shield (Zymo Research, Irvine, CA, USA) for subsequent DNA extraction.

### 2.2 In vitro batch fermentation (NCT/NFT dosing comparison)

Each enterotype was inoculated (1% v/v) into 1 mL of mBHI medium in a 96-well deep well plate or the same medium supplemented with soluble starch (0.35% w/v), methylcellulose (2% w/v), or NCT/NFT (2, 0.7, 0.5, 0.25, 0.1, 0.05, or 0.01% w/v), with six independent replicates per enterotype, per treatment. At inoculation, 100 µL of each replicate was transferred to a 96-well plate sealed with a sterile, gas-permeable sealing film (Diversified Biotech, Dedham, MA, USA) and incubated, as stated previously. The 96-well deep well plate was sealed with a sterile, breathable sealing film (Celltreat Scientific Products, Pepperell, MA, USA), allowing for gas exchange, and incubated, as discussed above. After 24 hours, the 96-well deep well plate was removed from the anaerobic chamber and centrifuged, as mentioned above. From each cell pellet 500 µL of cell-free supernatant was collected, transferred into a new 96-well deep well plate, and stored at -80°C. Cell pellets were resuspended in DNA stabilization buffer (Zymo Research, San Diego, CA, USA) for DNA extraction.

### 2.3 DNA extraction and 16S rRNA sequencing

DNA was extracted from pelleted cells as described previously^16^ using a ZymoBiomics DNA Miniprep kit (Zymo Research; Irvine, CA, USA) according to the manufacturer’s instructions. The protocol required each sample to undergo a cycle of bead-beating for one minute in an MP Biomedical FastPrep 24, followed by a one-minute incubation on ice for a total of five cycles. The DNA obtained underwent 16S rRNA V3/V4 amplicon sequencing using a dual-indexed barcoding strategy as previously outlined^17^, with adaptations to the amplification sequences^18,19^. A HEPA-filtered laminar flow cabinet intended for PCR preparation was used to generate the amplicons. Reactions were conducted by adding to 2 µl template DNA: 200 nM of each primer, 0.5 mM of MgCl_2_, and GoTaq Master Mix (Promega; Madison, WI, USA) for a total volume of 25 µl per sample. An MJ Research PTC-200 thermocycler was programmed as follows: one cycle of 94°C for 3 min; 94°C for 45 s, 50°C for 60 s, and 72°C for 90 s for a total of 25 cycles, and a final extension of 72°C for 10 min. PCR products were pooled and purified with a DNA Clean & Concentrator purification kit (Zymo Research, San Diego, CA, USA). The pooled amplicon library was sequenced on an Illumina MiSeq platform at the Idaho State University Molecular Research Core Facility (RRID:SCR_012598).

### 2.4 Sequencing data processing and analysis

*S*equencing data was demultiplexed and analyzed with QIIME 2™^20^. Reads were trimmed, joined, and denoised before assigning reads to amplicon sequence variants (ASVs) using DADA2^21^. Representative sequences were aligned with FastTree^22^, and taxonomic assignments were assigned using the Greengenes database (13_8, 99%) with a naïve Bayesian feature classifier trained against the representative sequences^20^. Differential abundance testing was completed using ANCOM-BC^23^, using a minimum prevalence cutoff of 0.001% and a Bonferroni correction for multiple comparison testing, with α < 0.05 considered as significant.

Sample rarefaction was performed to generate diversity measures, including the Shannon entropy^24^, as well as weighted UniFrac ^25^ and Bray Curtis ^26^ distance matrices. Inter-group differences in community composition were compared using PERMANOVA^27^ for inter-group comparisons and adonis^28^ for single-parameter or complex interactions from distance matrices. Continuous numerical metadata was transformed into a distance metric and compared to community composition using the Mantel test^29^. Sample enterotypes were determined from genus-level abundance data using the method available at https://enterotype.embl.de and previously described by Arumugan *et al*.^14^.

### 2.5 Gas Chromatography - Mass Spectrometry (GC-MS) analysis

Samples were thawed on ice and centrifuged at 4 ºC for five minutes at 14,000 rpm. 50 μL of supernatant aliquots were transferred into glass GC-MS vials with inserts. For GC-MS analysis, a Varian 3800 GC (Zebron ZB-FFAP, 30 m, 0.25 mm, 0.25 μm, Phenomenex, CA), 4000 Ion Trap MS (Varian) equipped with electron ionization source was employed. The vials were placed on the instrument tray in a randomized fashion. 1 μL was injected into the GC inlet with split ratio of 1:100; the syringe was washed with methanol to prevent carryover for the next injection. The ion trap was set to 250°C, the MS set to 100°C, and the transfer line set to 250°C. The GC protocol was set as follows: starting with 50°C and increasing 15°C per minute until 220°C, and a 2 min hold to purge the column. The helium carrier gas was set to a constant 2 mL/min flow. The m/z range scanned was 50–200 Th. Empty vial blanks and collected experimental blanks were randomly interspersed with the samples. The SCFA reference samples were injected in triplicates after the samples to preclude possible carryover. Collected GC-MS data were then analyzed offline. Free fatty acid (SCFAs) test mixture (Restek, Cat#35272, 1000 ug/mL) was diluted to 500, 250, 125, 62.5, 31.25, 15.625, 7.8125 and 3.90625 μg/mL concentrations to generate calibration curves. The raw data were converted from the vendor’s format to cdf. The deconvolution was carried out using MSHub with fully automatic settings, as described by Aksenov et al^30^. For chemical identification, the mass spectra were matched using the NIST 2020 MS library. The peaks for target SCFAs were manually identified using reference data for the standards. Concentration calculations were carried out by using formulas for each calibration curve for respective SCFAs.

### 2.7 Additional statistical methods

Statistical tests were performed in R (v. 4.2.2)^31^, using the nonparametric Kruskal-Wallis test^32^ for group comparisons and the Wilcoxon rank-sum test^33^ for individual comparisons using *ggpubr* (v. 0.4.0) and *rstatix* (v. 0.7.0) R packages^34,35^. Multiple comparisons included the Bonferroni correction for multiple comparison tests ^36^. Statistical correlations were calculated using a Spearman correlation^37^. Growth curve parameters were calculated using the *growthcurver* (v. 0.3.1) R package. Figures were visualized using the *tidyverse, ggplot2* (v. 3.4.0), *ggpubr* (v. 0.4.0), and *viridis* (v. 0.6.2) R packages^34,38,39,40,41,42^.

## 3. Results

### 3.1 Microbiomes shift in response to Bio Gut Fiber™ and NCT/NFT

From the microbial communities grown to 24 hours post-inoculation, recovered DNA was subjected to 16S rRNA amplicon sequencing, resulting in 15.8M paired-end reads passing quality filtering and demultiplexing. The mean number of read pairs per sample was 39,946, and the median number of read pairs per sample was 39,058 across a total of 115 samples. To calculate diversity measures, samples were rarefied to 20,000 reads per sample, omitting samples with fewer reads per sample.

In assessing the inter-sample variability (β diversity), a weighted UniFrac distance matrix explained the most variability across the samples (86.0%), and the Bray Curtis distance metric also performed well (75.5%), but both were substantially more than the unweighted UniFrac distance metric, which explained only 46.1% of the inter-sample variability. Using the weighted UniFrac distance metric, we found that both the starting inoculum (E1, E2, or E3) and treatment (starch, methylcellulose, Bio Gut Fiber™, or NCT/NFT) resulted in significant community composition differences when compared by an adonis test, as well as the interaction of both treatment and enterotype (P = 0.001 for all comparisons; Figure 1A-B).

**Figure 1.**
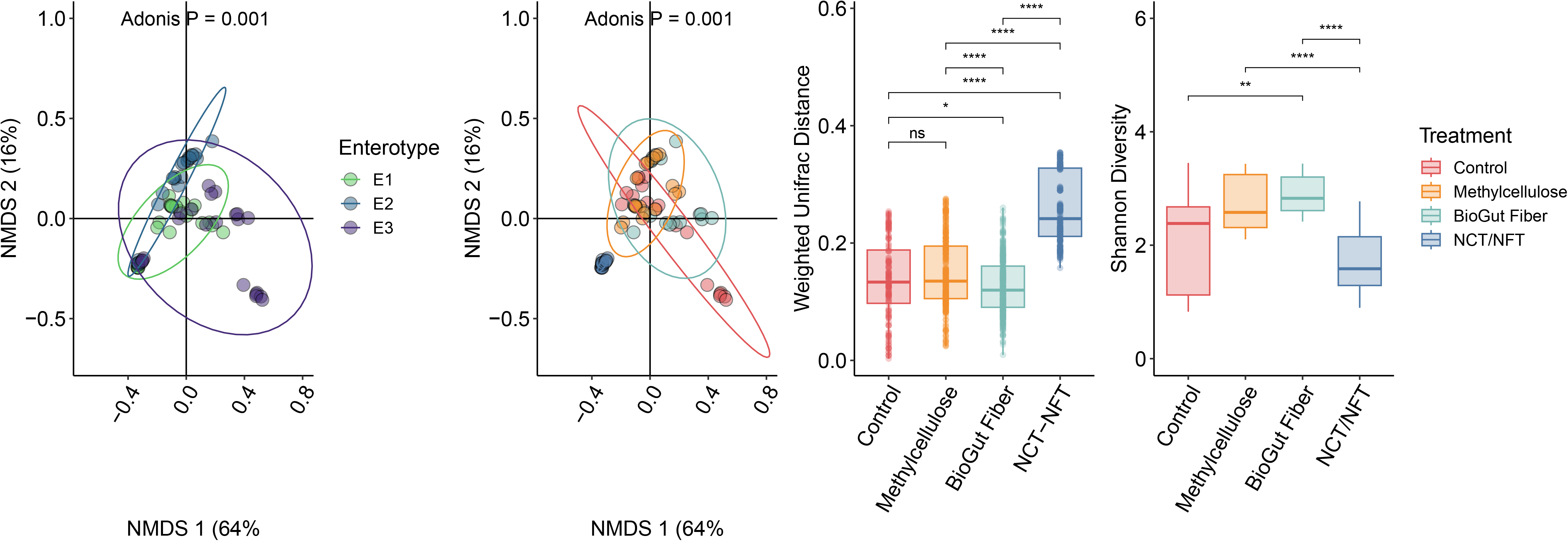
Community composition shifts in response to substrates provided across communities tested. A weighted UniFrac distance metric captured the greatest proportion of variation (80% in the first two axes) among the distance measures provided. These PCoA plots demonstrated a strong effect of both the inoculum (A) and the substrate (B) when compared using adonis test. The greatest degree of shift was observed among NCT/NFT-fed communities and (C) showed lower community diversity relative to the Bio Gut Fiber™fed communities (D). (ns, not significant; *, P < 0.05; **, P < 0.01; ****, P < 0.0001).

We then examined which groups were significantly different from the starch control and found that methylcellulose, NCT/NFT, and Bio Gut Fiber™ were significantly different when all groups were compared together using a PERMANOVA test on the weighted UniFrac distance measures. However, while we found that the methylcellulose, Bio Gut Fiber™ and NCT/NFT were all significantly different from the control within inocula E2 and E3 (P < 0.05 for E2, P < 0.01 for E3), inoculum E1 did not show a significant difference between Bio Gut Fiber™ and the control (P > 0.05), while the other comparisons were significantly different (P < 0.01).

In addition to the differences in community compositions, we also found that across inocula, the community richness and evenness, Shannon entropy, was significantly higher among the methylcellulose and Bio Gut Fiber™-treated communities, while the NCT/NFT-treated communities declined in community diversity relative to the control communities (Figure 1C-D). When each inoculum was considered individually, we found that Bio Gut Fiber™-treated communities were significantly enriched in terms of diversity relative to the starch control medium, and while community diversity in the methylcellulose-treated inocula was significantly higher than the starch control medium among inocula E2 and E3, community diversity was lower than the starch control medium in community E1. Across E1 and E2 communities, NCT/NFT-treated communities were significantly lower in terms of community diversity than the starch control medium, while the E3 community increased in diversity relative to the starch control medium (P < 0.001, Wilcoxon test).

To determine whether the changes in community diversity and differences in community composition were related to community productivity (i.e., growth), we used measurements of the optical density at 600 nm (OD_600nm_) to monitor cell density throughout the 24-hour growth period. Using these growth curves, we determined the area under the curve (AUC) to compare growth curves across enterotypes and treatments. As the AUC is a function of both time and maximum cell density, we reasoned that this was a useful metric to examine growth as we observed diauxic growth characteristics in some of the conditions tested (Figure 2A-C). When we compared the AUC across treatments, we found that communities incubated with Bio Gut Fiber™ exhibited greater AUC compared to the starch control or methylcellulose-fed communities (Figure 2D). Comparably, communities grown on Bio Gut Fiber™ or NCT/NFT showed the most rapid time to their midpoint in growth, suggesting a more rapid growth phenotype overall (Figure 2E).

**Figure 2.**
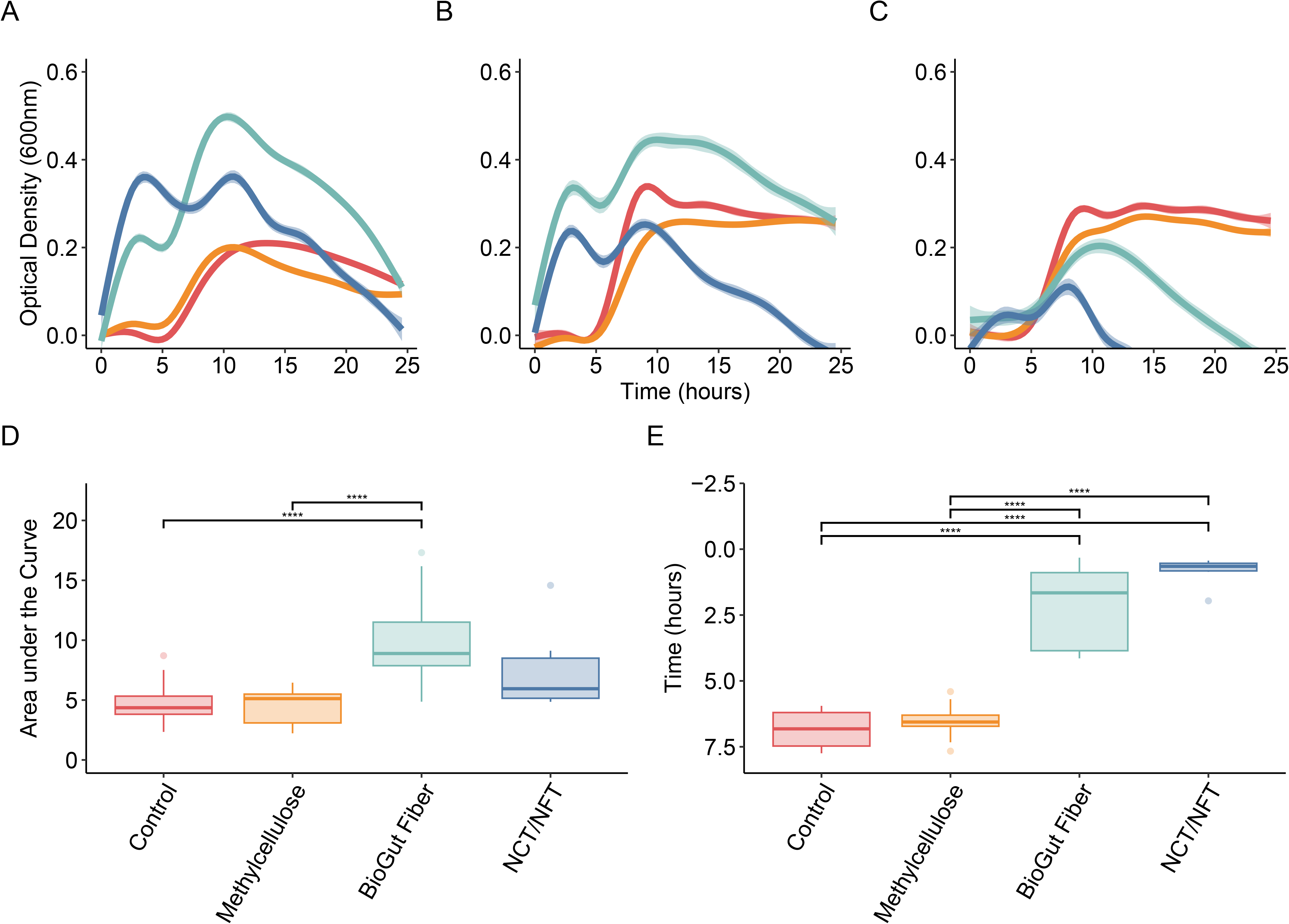
Growth kinetics differed across communities tested. Inoculum E1 (A), E2 (B), and E3 (C) showed distinct growth kinetics over 24 hours when the optical density at 600 nm (OD_600nm_) was measured. In both E1 and E2 (A and B), a diauxic-type curve was observed among communities incubated with Bio Gut Fiber™, with a pause around 5 hours of growth. When these growth curves were evaluated, (A) the area under the curve (AUC) and (E) the time to mid-point in growth, were strongest among Bio Gut Fiber™-fed communities, while the control and methylcellulose growth kinetics were weaker. NCT/NFT-fed communities demonstrated faster time to the growth midpoint (E) but had a weaker maximum cell density (A-D). (ns, not significant; *, P < 0.05; **, P < 0.01; ****, P < 0.0001).

### 3.2 Family-level taxonomic changes

As the growth phenotypes of the inoculated communities, as well as their community-level and sample-level diversity measurements indicated strong phenotypic differences, we examined the taxa predominant among these communities. Notably, the greatest enrichment observed was for *Enterobacteriaceae* among the NCT/NFT-fed communities, with this family becoming the predominant bacterial group after 24 hours (Figure 3A-C).

**Figure 3.**
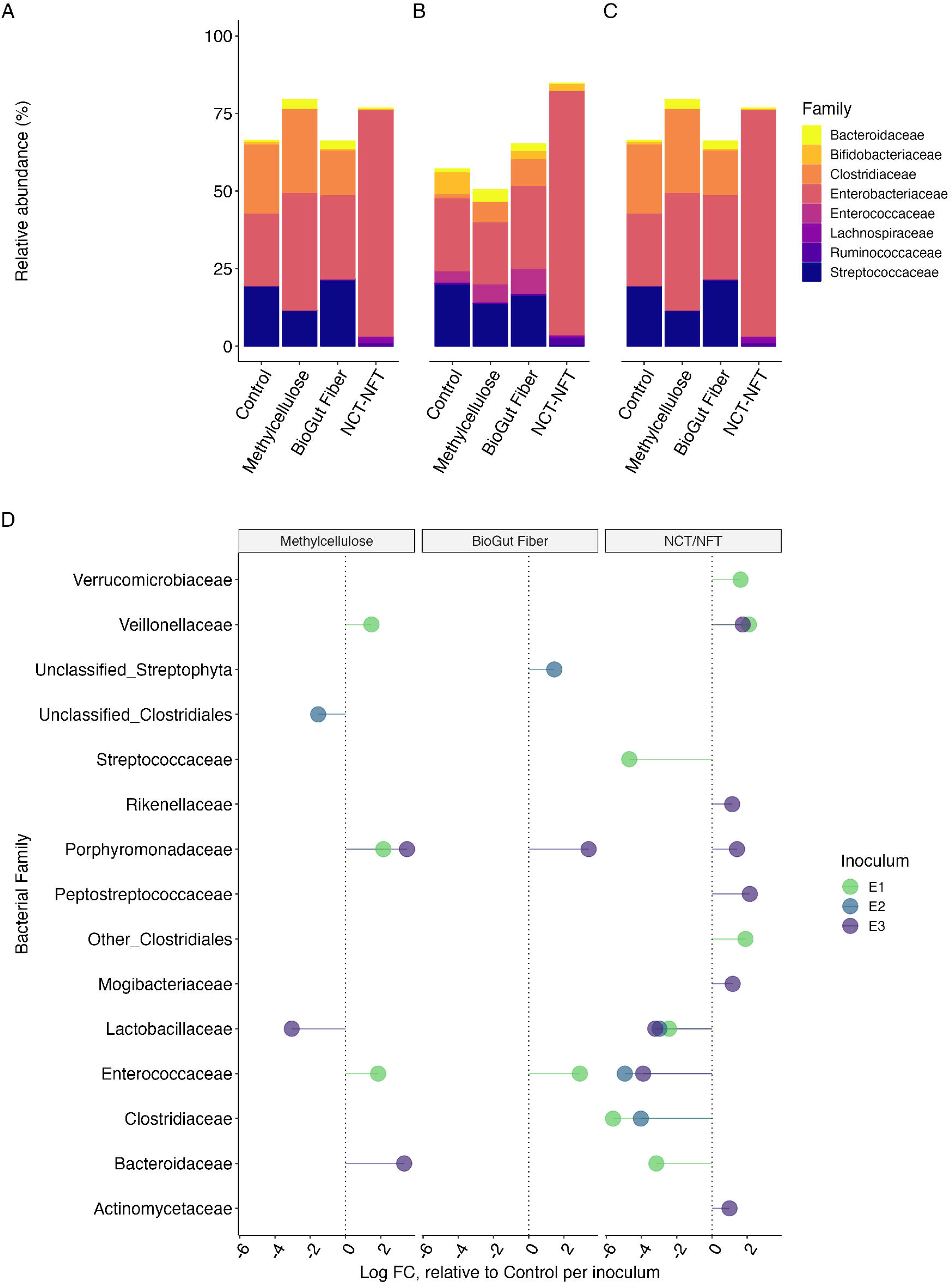
Community compositions differ most strongly among the NCT/NFT-fed communities. Across the three inocula, E1 (A), E2 (B), and E3 (C), NCT/NFT-fed communities exhibited a strong dominance of *Enterobacteriaceae*, relative to the other treatments across inocula. Bio Gut Fiber™-fed communities more closely resembled the control and methylcellulose-fed communities but with more rapid growth (see Figure 2). D) ANCOM-BC was used to identify differentially abundant bacterial families across the treatments and inocula. The distance from the central dotted line (at X=0) indicates the log fold change relative to the control communities for each inoculum (points are colored by inocula). All comparisons shown are statistically significant after Bonferroni correction (q < 0.05).

To identify differentially abundant taxa, we used ANCOM-BC to examine the effect of inoculum and substrate on the final community composition. To compare effects across treatments, we examined the differences of the substrates relative to the control media that was not supplemented for each of the inocula tested. Relative to the control, communities incubated with methylcellulose showed increased relative abundance of *Veillonellaceae, Porphyromonadaceae*, and *Enterococcaceae* (q < 0.05) among the E1-inoculated communities, Unclassified *Clostridiales* were diminished among the E2-inoculated communities, relative to the control (q < 0.05), and *Porphyromonadaceae* and *Bacteroidaceae* were increased among the E3-inoculated communities, relative to the control, while *Lactobacillaceae* were decreased (q < 0.05). Bio Gut Fiber™ increased *Enterococcaceae* among the E1-inoculated communities, apparent *Streptophyta* (possible chloroplast remnants from the Bio Gut Fiber™) in the E2-inoculated communities, and *Porphyromonadaceae* among the E3-inoculated inoculum (q < 0.05 for all comparisons). Among communities treated with 2% (w/v) NCT/NFT, *Verrucomicrobiaceae, Veillonellaceae*, and unclassified *Clostridiales* were increased among the E1-inoculated communities, while *Veillonellaceae, Rikenellaceae, Porphyromonadaceae, Peptostreptococcaceae, Mogibacteriaceae*, and *Actinomycetaceae* were increased among the E3-inoculated community, relative to the respective control communities (q < 0.05). Among the NCT/NFT-treated communities, *Streptococcaceae, Lactobacillaceae, Clostridiaceae*, and *Bacteroidaceae* were decreased among the E1-inoculated communities; *Enterococcaceae, Lactobacillaceae*, and *Clostridiaceae* were decreased among the E2-inoculated communities; and *Lactobacillaceae* and *Enterococcaceae* were decreased relative to the control among the E3-inoculated communities (Figure 3D).

### 3.3 Short-chain fatty acid production

Quantification of SCFA production identified significant differences across the control, methylcellulose, and Bio Gut Fiber™ groups. Acetate, butyrate, and propionate were all significantly different across these groups when compared by a Kruskal Wallis test (P < 0.05; Figure 5). In particular, an *ad hoc* Wilcox test indicated that acetate was higher among Bio Gut Fiber™ treated communities (9584 μg/mL +/-2934 μg/mL SD), relative to the methylcellulose control (5806 μg/mL +/-1597 μg/mL SD; FDR-adjusted P < 0.01), while butyrate was higher among the Bio Gut Fiber™ (658 μg/mL +/-392 μg/mL SD) and methylcellulose-treated communities (347 μg/mL +/-282 μg/mL SD), relative to the control (66 μg/mL +/-105 μg/mL; FDR-adjusted P < 0.05). Finally, propionate was significantly higher among the methylcellulose (502 μg/mL +/-396 μg/mL SD) and Bio Gut Fiber-treated communities (709 μg/mL +/-374 μg/mL SD) relative to the control (97 μg/mL +/-67 μg/mL SD; FDR-adjusted P < 0.001; Figure 5).

### 3.4 The effect of NCT/NFT is dose-dependent on in vitro communities

As the differences observed among communities grown on NCT/NFT were quite stark (Figure 1-3), we next examined the impact of the concentration of NCT/NFT to understand if these differences were an effect of the concentration of these tyramines on the community composition and phenotype. Using the same inocula, we prepared a second 24-hour in vitro fermentation using NCT/NFT concentrations of 2%, 0.7%, 0.5%, 0.25%, 0.1%, 0.05%, and 0.01% (w/v), along with the same control groups (2% starch and 2% methylcellulose, w/v).

After performing sequencing on these samples, 15.9M read pairs passed quality filtering with an average of 88,042 read pairs per sample, ranging from 26,704 to 170,175 read pairs per sample. After processing these data in the same manner (See Methods), we found a strong correlation between community composition, assessed by a weighted UniFrac distance metric, and the NCT/NFT concentration (P = 0.001, rho = 0.84; Mantel Test). This association was remarkably strong across all communities (rho = 0.51, 0.85, and 0.83 for E1, E2, and E3, respectively; P = 0.001 for all comparisons).

The effect of this association was apparent in the weighted UniFrac PCoA (Figure 4A-B) and was significantly correlated with the abundance of *Enterobacteriaceae* (rho = 0.78, 0.87, and 0.74, P < 0.001) across all three inocula (Figure 4C).

**Figure 4.**
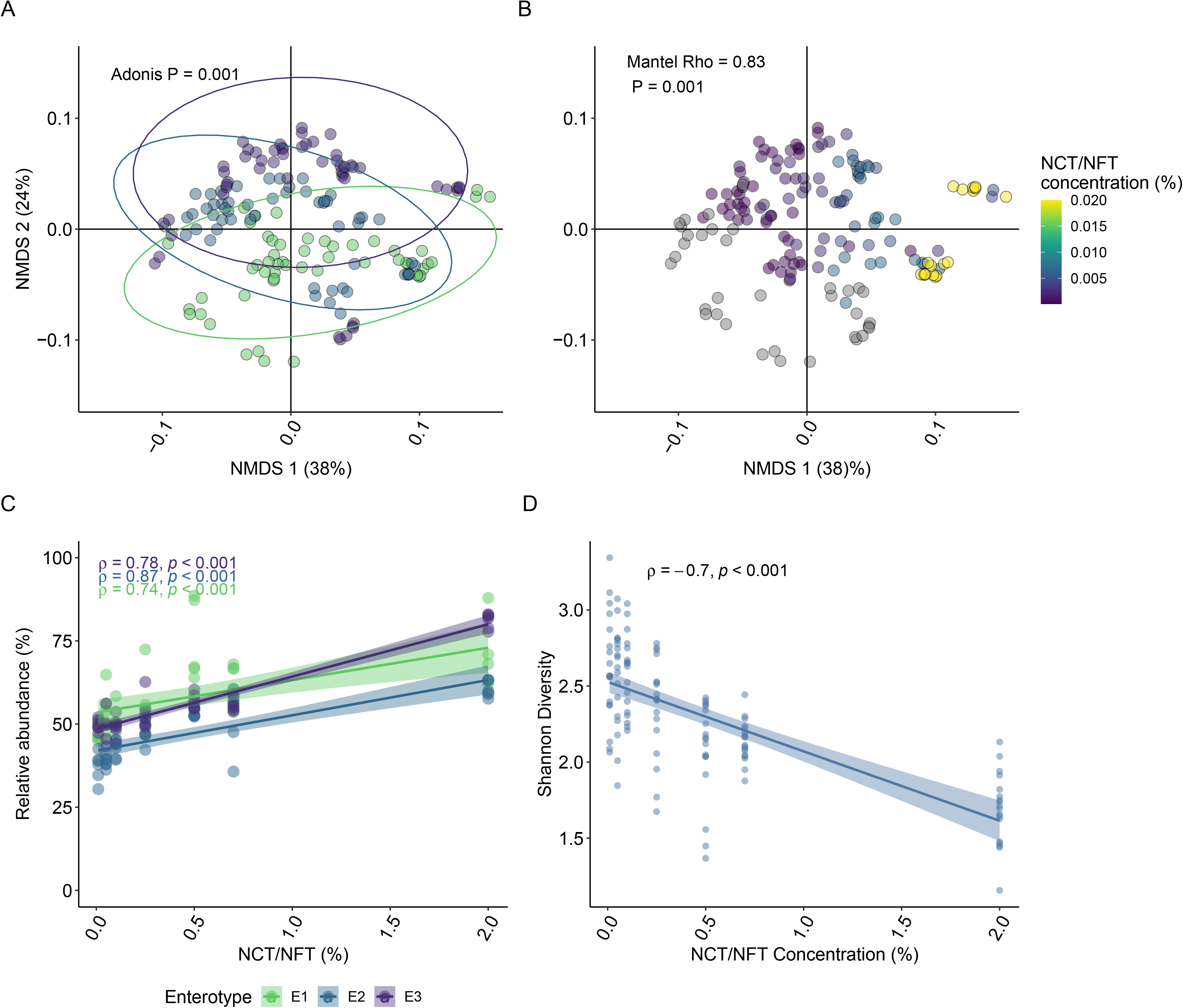
The concentration of NCT/NFT drove shifts in the community composition. Among inocula treated with a range of NCT/NFT concentrations (w/v), the inocula-specific differences observed previously were again confirmed (A), while the NCT/NFT concentration strongly correlated with the community distance metric (B) by a Mantel test (rho = 0.83, P = 0.001). The abundance of Enterobacteriaceae was also strongly associated with the NCT/NFT concentration (C) as well as the decline in community diversity (D).

## 4. Discussion

Considerable interest has been focused on understanding how secondary plant compounds such as polyphenols or fiber affect the gut microbiome and researchers have been particularly interested in leveraging these interactions to improve human health.^2, 33, 34^ Despite this interest, little is yet known about how polyphenols and fiber impact the gut microbiome and may ultimately affect host health. Instead, researchers often examine correlational data estimated from dietary intake as assessed through a dietary recall or food frequency questionnaire.^35^ In parallel, significant progress has been achieved in understanding how specific structural features of key fibers affect the gut microbiome, with a major interest in classes of oligosaccharides derived from human milk as prebiotic substrates.^36^ Through the isolation of specific structures, this area of research has been able to build extensive mechanistic connections between the introduction of particular fiber structures (i.e. human milk oligosaccharides, HMOs) and the impact that these structures have on the gut microbiome, how genetic loci enable key taxa to thrive in the infant gut microbiome when fed HMOs, and how they may be leveraged to improve infant formula.^37-39^

To begin to bridge this gap in knowledge among plant-derived bioactives, we performed a combination of *in vitro* fermentation and 16S rRNA sequencing to assess the effect of a milled hemp hull, Bio Gut Fiber ™, containing ∼0.7% w/v NCT/NFT and extracted NCT/NFT in more comparable proportions as found in Bio Gut Fiber ™ on model human gut microbiomes *in vitro*. To provide a relevant comparison, we added a starch control medium and methylcellulose, often sold as a fiber supplement but structurally similar to cellulose found in Bio Gut Fiber™, though lacking the same polyphenolic and bioactive compounds found within the hemp hull food matrix. Here, we used three pooled human gut microbiome inocula that vary in functional composition and species,^10^ providing a repeatable and tractable model of microbiome responses across humans generally.

We found that the introduction of Bio Gut Fiber™ and NCT/NFT significantly altered individual communities relative to the methylcellulose or starch controls (Figure 1-3), but also found that the addition of the hemp hull product, Bio Gut Fiber™, generated unique responses relative to the NCT/NFT ingredient. Despite otherwise comparable increases in community productivity (Figure 3), we found that community diversity increased the most, relative to the starch control, among the communities fed Bio Gut Fiber™ as a substrate (Figure 1). While community diversity is often reported as a key component of gut microbiome ‘health’, the role of community diversity is more nuanced.^40^ Among macroecology systems, community productivity is an important measure of ecosystem fitness, with greater productivity generally representing a thriving community, though not always correlative with alpha diversity.^41^ Moreover, short chain fatty acid (SCFA) production had the greatest mean concentration among communities treated with Bio Gut Fiber™ across acetate, propionate, and butyrate (Figure 5), which may be another measure of a productive community and can be a beneficial product in the gut microbiome.^7^

**Figure 5.**
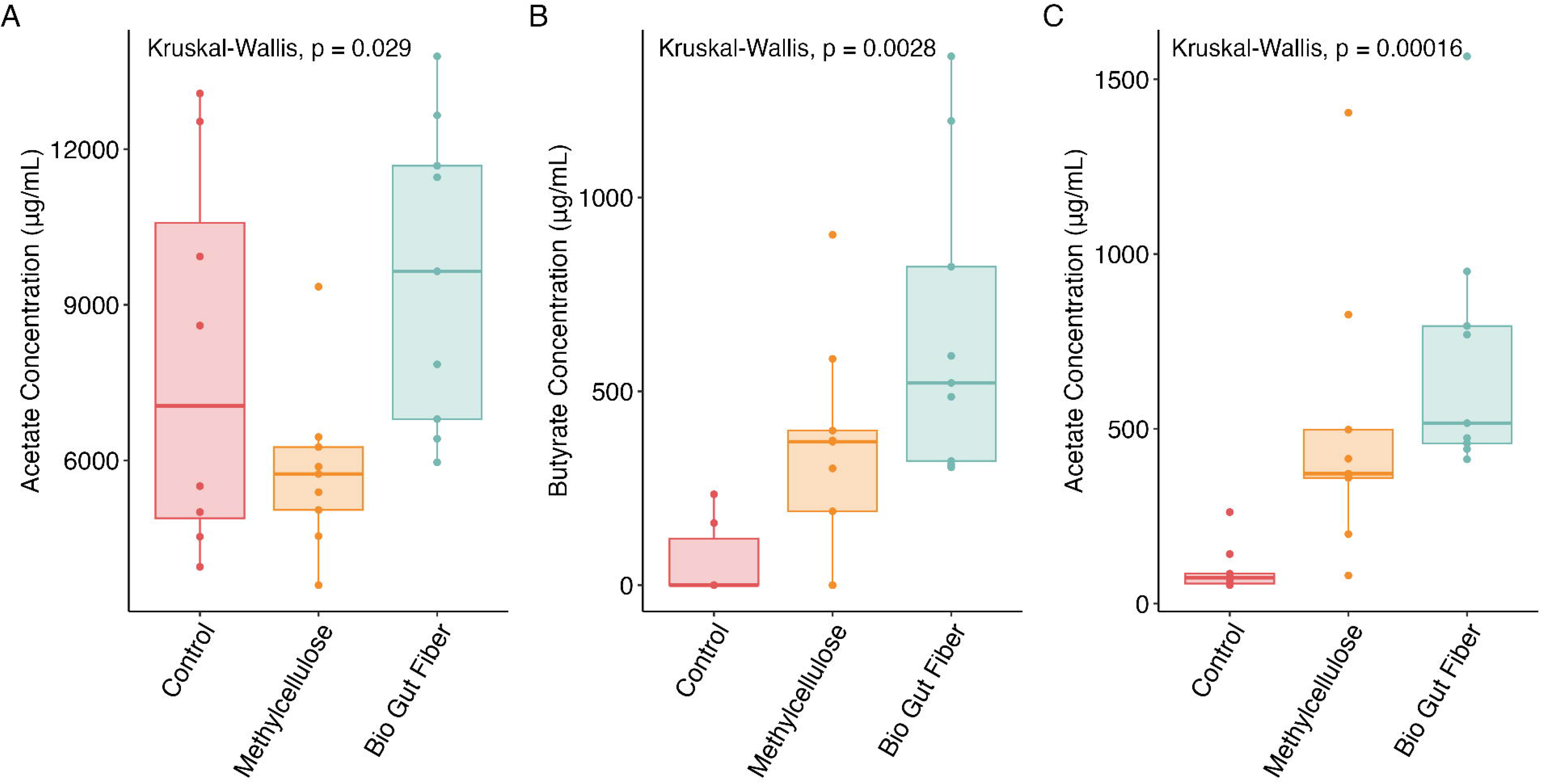
Short-chain fatty acid concentrations trend higher in Bio Gut Fiber™ samples. Quantitation of acetate (A), butyrate (B) and propanoate (C) were significantly different across the three tested groups (control, methylcellulose, and Bio Gut Fiber™). Acetate concentrations were significantly higher in the Bio Gut Fiber™ relative to the methylcellulose-treated communities (P < 0.05), while butyrate and propanoate concentrations were higher in Bio Gut Fiber™ treated communities relative to the control samples, but not methylcellulose treated communities.

Indeed, here we observe deficits in terms of community (alpha) diversity among the samples incubated with high concentrations of NCT/NFT but find that those changes are also linked to diminished community productivity (Figure 3D), even though the rate of community growth (Figure 3E) is unchanged. Importantly, the differences in community diversity in this model are a product of an increase in the proportion of *Enterobacteriaceae*, and while enterobacteria are associated with enteric inflammation,^42,43^ our second experiment demonstrated that the increases in *Enterobacteriaceae* observed initially are likely a result of the initial concentration of NCT/NFT (2% w/v). The 2% concentration is substantially higher than in Bio Gut Fiber™ (∼0.7% w/v) or what could be encountered *in vivo*. When we compared the lower doses comparable to what is observed in the Bio Gut Fiber™ product, the levels of *Enterobacteriaceae* were no longer enriched, and the overall community composition was more similar to the methylcellulose and starch controls (Figure 4).

At present, the mechanism by which NCT and NFT may affect the gut microbiome remains elusive. Most work to date has been conducted with hemp seeds which are rich in omega-3 fatty acids^44^ or hemp seed bran,^45^ which is also distinct from hemp seed hulls. Together, our findings show that an *in vitro* model of the human gut microbiome finds microbiome-specific trends in community composition, community productivity, and the magnitude of the effect of Bio Gut Fiber™ or NCT and NFT on human gut microbiomes *in vitro*. Additionally, this approach isolates the effect of these ingredients on the gut microbiome from host factors which could mitigate or influence the impact on the gut microbiome, which is both an advantage and a limitation of the current work. Additional work will be needed to understand how Bio Gut Fiber™, NCT, and NFT affect the gut microbiome *in vivo*, and whether we see similar effects across individuals whose gut microbiomes are compositionally and functionally similar to each of the three inocula tested here.

## 5. Data availability

Sequencing data from the experiments described here are available in the NCBI SRA under BioProject PRJNA1104297.

## 6. Acknowledgements

The authors thank the Molecular Research Core Facility at Idaho State University, RRID:SCR_012598, for conducting the Illumina sequencing used in this study.

